# Generation of an optimised Cumate toolkit for tuneable protein expression during *in vitro* and *in vivo* studies of *Burkholderia cenocepacia*

**DOI:** 10.1101/2025.11.26.690688

**Authors:** Hamza Tahir, Kristian I. Karlic, Godfrey Mwiti, Nichollas E. Scott

## Abstract

Inducible gene expression is pivotal for dissecting bacterial physiology and virulence mechanisms. Across the *Burkholderia* genera, a limited range of inducible systems currently exist that show minimal impacts on the proteome and allow tight regulation. In this study, we engineer a set of cumate inducible vectors for use in *Burkholderia cenocepacia* that offer minimal basal expression and the ability to control *B. cenocepacia* gene expression within Eukaryotic cells. Through mutagenesis-based studies of cumate circuits and the cumate regulator (CymR), we generate an optimized cumate circuit (P_CymRC_/CymR_GV_) which allows the tight and tunable control of protein expression within *B. cenocepacia,* as assessed by fluorescent and protein O-linked glycosylation analysis. Using comparative proteomics, we demonstrate cumate induction leads to both reduced and orthogonal effects on *B. cenocepacia* compared to widely used rhamnose based induction systems. Leveraging the cell permeability of cumate and the generation of a CTX-based chromosomal integration vector, we show that inducible control of protein expression is achievable during intracellular replication of *B. cenocepacia.* Finally, using the ability to control intracellular expression, we demonstrate the requirement of O-linked protein glycosylation for optimal *B. cenocepacia* intracellular replication. Combined, this work demonstrates that cumate inducible systems allow precise and tuneable gene expression in *Burkholderia* even within a host-pathogen context.

**Importance:** This work establishes optimised cumate-inducible vectors for use in *Burkholderia cenocepacia*, addressing the need for alternative inducers to available carbohydrate systems. We show cumate-inducible vectors allow precise control of gene expression even within eukaryotic cells, providing a new and orthogonal way to temporally control protein induction. Utilising cumate- based induction, we demonstrate the importance of O-linked protein glycosylation for optimal intracellular replication in *B. cenocepacia*, highlighting its potential to be used to explore host-pathogen interactions. Combined, this work shows cumate-inducible vectors extend the range of studies which can be undertaken to dissect *B. cenocepacia* physiology and virulence.

## Introduction

Precise, titratable modulation of gene regulation is indispensable for studying bacterial physiology and virulence at both the mechanistic as well as systems levels [1–3]. Ideally, inducible systems should provide rapid response kinetics, high dynamic range and minimal basal expression to allow the precise assessment of gene functions in a controllable manner [4–6]. Over the last 50 years, a diverse array of inducible systems responsive to naturally occurring as well as synthetic or non-metabolizable inducible agents have been developed [5, 7–10]. Within commonly utilised inducible systems the control of gene expression is typically mediated by regulatory circuits composed of promoters and repressor pairs with well-known examples including P_lac_/LacI [11], P_tet_/TetR [12], P_BAD_/AraC and P_rha_/RhaS [13] systems. To improve the control of gene expression, synthetic biology, including protein engineering as well as rational DNA regulatory designs are increasingly utilized to generate refined regulatory circuits [3, 9, 10]. To date several engineered regulatory circuits have been reported with a key goal of these systems being to both reduce basal expression and enhance specificity of circuits compared to the canonical/natural regulatory systems [3, 14, 15]. While early studies focused on well-known regulatory circuits to achieve these goals [16] recent efforts have sought to expand the toolbox of engineered regulatory systems by refining novel circuits for diverse applications such as metabolic engineering [17] with one such regulatory circuit being the 4-isopropylbenzoate inducible circuit.

4-isopropylbenzoate, here in referred to as *c*umate, is the native ligand for the P_Cym_/CymR inducible circuit identified within the *cmt* operon of *Pseudomonas putida* F1 [18–20]. Within this system, repression of cumate catabolism is mediated by the binding of CymR to a <u>Cu</u>mate <u>O</u>perator (CuO) DNA sequence which in the presence of cumate dissociates, allowing gene expression downstream of CuO [20]. Due to cumate being cell permeable, nontoxic as well as non-metabolizable in the absence of a cumate metabolism pathway [19, 20], several teams have used the canonical P_Cym_/CymR inducible circuit for titrable gene expression within *Escherichia coli* [21, 22], *Pseudomonas aeruginosa* [23] as well as non-model organisms such as *Streptomyces spp* [24] and members of the *Alphaproteobacteria* [25]. Notably, the high permeability and non-toxic nature of cumate makes this inducer an attractive option within systems where other agents may be poorly tolerated or excluded which is best exemplified by its use in mammalian systems where eukaryotic cumate-inducible circuits have been constructed [26]. In 2018 Meyer *et al* engineered a synthetic cumate circuit, referred here in as P_CymRC_/CymR_AM_, which has been suggested to possess reduced basal expression and minimal antagonism with other commonly used inducers [3]. Due to the accessibility of this engineered variant several teams have incorporated P_CymRC_/CymR_AM_ within plasmids designed for non-model organisms and demonstrated cumate induction of fluorescent reporters [27, 28]. However, as this engineered regulatory circuit has only been assessed within a limited context to date, it is unclear if its reported characteristics are a true reflection of its performance, especially within non-model organisms such as members of the *Burkholderia* genera.

*Burkholderia* species are ubiquitous environmental organisms, and while the majority are non-pathogenic, members of *the Burkholderia cepacia* and *Burkholderia pseudomallei* complexes are associated with serious human infections [29–31]. Within the *Burkholderia cepacia* complex, *Burkholderia cenocepacia* is a significant opportunistic pathogen in patients with cystic fibrosis (CF), where it is able to infect and persist intracellularly within host cells, driving a progressive decline in lung function [31, 32]. To date, the study of *B. cenocepacia* pathogenesis has been impeded due to both the innate multidrug resistance of isolates, which has limited the resistance markers applicable for the creation of gene knockouts [33, 34], as well as a paucity of tools for precise complementation and/or protein overexpression studies. Within *B. cenocepacia*, inducible protein expression studies have largely utilized two carbohydrate-based inducer systems, P_BAD_/AraC [35] and P_rha_/RhaS [36], activated by arabinose and rhamnose respectively. While these circuits are functional in a range of *Burkholderia* species, P_BAD_/AraC is known to display basal expression [37] and requires high concentrations (2–3%) for the induction of uniform protein expression [36, 38]. In contrast, P_rha_/RhaS has been shown to provide tight and uniform regulation of gene expression [36], making it the preferred induction system and leading to its use in several recently developed CRISPR systems [39, 40] as well as the assessment of essential genes [41, 42]. However, within our lab we have previously observed that the induction of the P_rha_/RhaS-based system with even modest rhamnose levels (0.1%) can evoke large proteomic changes within *B. cenocepacia* [43]. While teams have made recent improvements to the dynamic range of P_rha_/RhaS for the control of essential genes within *B. cenocepacia* [42], orthogonal chemically inert inducers with minimal proteomic effects would be highly beneficial for the control of protein expression within *Burkholderia* species.

Within this work we have sought to establish a set of cumate-inducible vectors to allow the tight regulation of protein expression within *B. cenocepacia*. Comparing available cumate regulatory circuits, using both fluorescent reporter and an enzymatic reporter system assessing protein O-linked glycosylation, we demonstrate that currently available circuits show high basal expression, which limits their utility for the study of enzymatic processes. Using targeted mutagenesis, we show that the previously reported point mutations suggested to enhance CymR_AM_ performance increases basal expression as well as potentiate the responsiveness to cumate. By combining elements of the canonical and engineered CymR_AM_ cumate circuit we have generated a new variant, P_CymRC_/CymR_GV_ that markedly reduces basal expression while preserving robust, dose-dependent induction. Deploying P_CymRC_/CymR_GV_ to control protein glycosylation, this circuit demonstrates precise, inducer-dependent modulation, with quantitative proteomics also revealing minimal perturbations following cumate induction. Finally, we show the use of cumate based induction provides a novel means to control protein expression within infection contexts. Collectively, this study establishes a novel set of cumate-inducible vectors for *B. cenocepacia,* as well as other *Burkholderia* species, that are orthogonal to current systems and are suitable to control protein expression within *in vivo* studies.

## Results

### The cumate circuit P_Cym_/CymR is functional yet displays high basal expression within *B. cenocepacia*

To expand the available protein expression vectors for *B. cenocepacia*, we sought to establish a set of cumate inducible systems and assess their performance. To allow the comparison of available cumate regulatory systems we first integrated the canonical circuit P_Cym_/CymR into the broad-host pBBR1 based vector pMLBAD [35], generating the plasmid pCumate-sfGFP (**Figure 1A & D**). Within *B. cenocepacia* K56-2, we assessed expression from pCumate-sfGFP revealing the dose dependent production of sfGFP as assessed by western blotting (**Figure 2A**). While fluorescent reporter proteins provide a convenient method to assess protein induction these systems can mask basal expression, which may occur below the dynamic range detectible by western blotting or direct fluorescent assessment. Based on this, we reasoned that the use of an enzymatic reporter may provide a more sensitive read-out for basal activity and placed the *B. cenocepacia* protein glycosylation oligosaccharyltransferase PglL (BCAL0960) [44] under cumate inducible control, generating pCumate-PglL*_Bc_*-his and assessed the restoration of glycosylation within the *B. cenocepacia* strain Δ*pglL* BCAL1086-his [45]. Within this background the periplasmic glycoprotein BCAL1086 has been chromosomally his-tagged, with prior studies demonstrating the loss of this protein in the absence of glycosylation, providing a sensitive readout for PglL activity [43, 45]. Introduction of pCumate-PglL*_Bc_*-his into Δ*pglL* BCAL1086-his resulted in the appearance of BCAL1086-his in the absence of induction, with the addition of cumate resulting in a decrease in gel mobility consistent with inducer-dependent glycosylation (**Figure 2B**). Glycoproteomic analysis confirms the introduction of pCumate-PglL*_Bc_*-his within Δ*pglL* BCAL1086-his partially restored glycosylation in the absence of induction (**Figure 2C, Supplementary Table 5)**. Consistent with this, quantitative analysis supports the presence of pCumate-PglL*_Bc_*-his is sufficient to result in the detection of several *B. cenocepacia* glycopeptides with glycosylation levels enhanced upon induction (**Figure 2D, Supplementary Table 6**). Taken together these results support that while P_Cym_/CymR is functional within *B. cenocepacia,* it displays high basal expression, as demonstrated by the detection of protein glycosylation in the absence of induction within a glycosylation-null complemented background.

**Figure 1.**
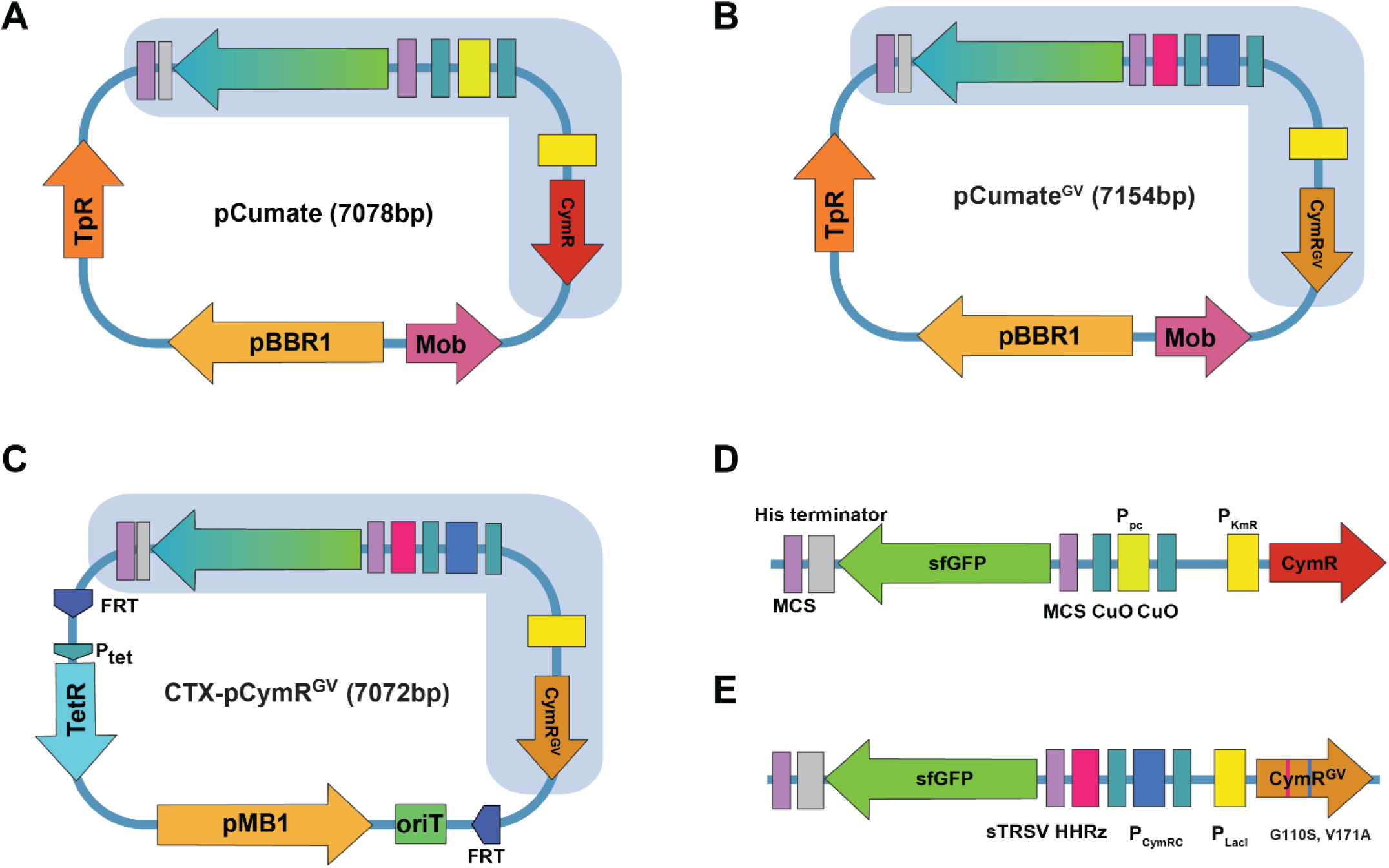
Schematic representation of cumate-inducible vectors and cumate circuit designs. **A)** Plasmid pCumate containing the P_pc_/CymR regulatory circuit within a pBBR1 backbone with trimethoprim resistance (Tp^R^) cassette and mobilization (Mob) elements. **B)** The optimised expression construct pCumateᴳⱽ, incorporating CymRᴳⱽ (G110S, V171A) for enhanced regulation and reduced basal expression. **C)** The CTX-pCymRᴳⱽ integrative vector utilising a Mini-CTX1 pMB1 backbone, FRT sites for flippase based marker excision and tetracycline resistance (tetR) cassette. **D–E)** Comparative schematic of cumate circuits, illustrating P_pc_/CymR and P_CymRC_/CymR_GV_ promoter configurations highlighting key regulatory elements including multiple cloning sites (MCS), Cumate operators (CuO) and Hammerhead Ribozyme (HHRz) components of the CymR and CymR_GV_ control systems.

**Figure 2.**
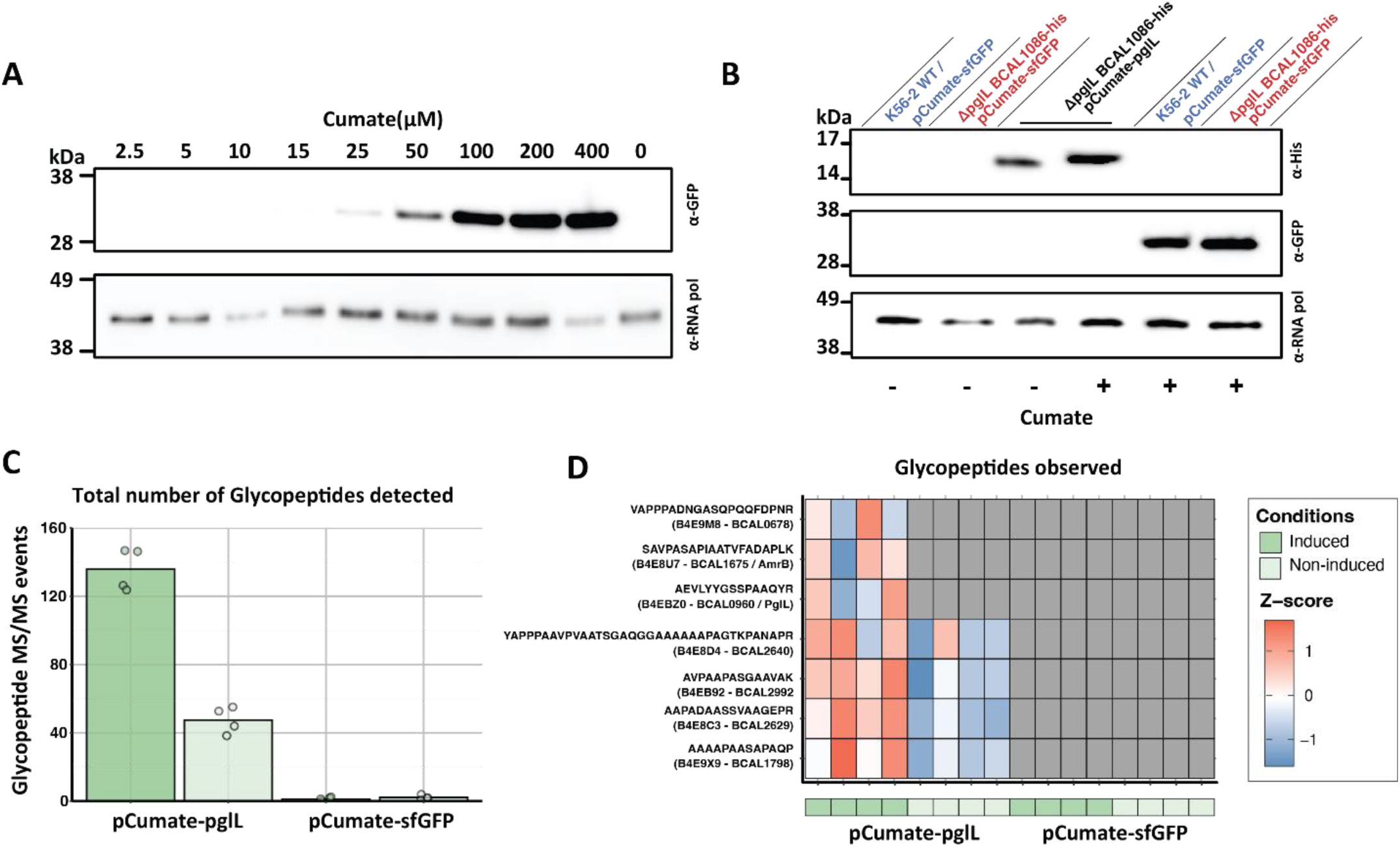
The cumate circuit P_Cym_/CymR displays high basal expression within *B. cenocepacia*. **A)** Western blot analysis demonstrates tuneable expression of sfGFP from pCumate-sfGFP in response to cumate. **B)** Immunoblot analysis of BCAL1086-his and sfGFP expression within the Δ*pglL* BCAL1086-his reporter strain revealing glycosylation in the absence of induction in a pCumate-PgIL_Bc_-his dependent manner. **C)** Quantification of glycopeptides identified from whole cell proteomic analysis of *B. cenocepacia* Δ*pglL* BCAL1086-his containing pCumate-PglL*_Bc_*- his or pCumate-sfGFP with and without induction with 100 μM cumate (n = 4). **D)** Heatmap of selected glycopeptides observed altered within *B. cenocepacia* Δ*pglL* BCAL1086-his reveals glycosylation in the absence of induction which is enhanced upon cumate induction (n = 4 per group).

### P_CymRC_/CymR_AM_ displays elevated basal expression compared to P_Cym_/CymR yet combining features of these circuits reduces basal expression

Our observation of high basal expression from P_Cym_/CymR suggests that to allow tight cumate- based control within *B. cenocepacia* the use of alternative cumate circuits may be required. As previous studies have demonstrated the functionality of P_CymRC_/CymR_AM_ within *Burkholderia thailandensis* [27] we assessed the performance of this circuit within *B. cenocepacia*. To allow a fair comparison of these circuits we exchanged P_Cym_/CymR within pCumate-sfGFP with P_CymRC_/CymR_AM_ from pKCyR5 [27], generating pCumate^AM^-sfGFP (**Figure 3A**). Surprisingly, we observed an increase in sfGFP expression when placed under P_CymRC_/CymR_AM_ control, compared to P_Cym_/CymR, in the absence of induction within *B. cenocepacia* (**Figure 3B).** We initially reasoned that expression changes due to codon usage within CymR_AM_, which was codon optimised for *E. coli* [3], may account for this increased expression. To assess this, we reintroduced the two amino acid substitutions from CymR_AM_, Ser110Gly (S110G) and Ala171Val (A171V) into the canonical CymR sequence within P_Cym_/CymR generating CymR^S110G^, CymR^A171V^ and CymR^S110G,A171V^. Introduction of these two mutations either individually or together resulted in increased basal sfGFP expression with the alteration of position A171V leading to a marked increase in expression which was partially restored by co-introduction of S110G **(Figure 3C).** These trends were also observed within *E. coli* (**Supplementary Figure 1A & B**) suggesting the prior reported alterations within CymR_AM_ are associated with unexpected impacts in the absence of induction. To confirm this hypothesis, we reverted the mutations within CymR_AM_ to the canonical amino acids, generating CymR_AM_^G110S^, CymR_AM_^V171A^ and CymR_AM_^G110S,V171A^. As within CymR, the alteration of position 171 led to a marked increase in basal expression which was suppressed by the alteration of position 110 (**Figure 3D**). Assessment of CymR_AM_, CymR and CymR_AM_^G110S,V171A^ revealed CymR_AM_^G110S,V171A^ resulted in reduced basal expression across the assessed cumate circuits (**Figure 3E**) with this reduction also observed within *E. coli* (**Supplementary Figure 1C**). These results support that CymR_AM_^G110S,V171A^, herein referred to as CymR_GV_ (**Figure 1B & E**), in conjunction with the promoter P_CymRC_ reduces basal expression, as assessed by sfGFP production, compared to P_Cym_/CymR and P_CymRC_/CymR_AM_ circuits within *B. cenocepacia*. Finally, to assess sfGFP expression in an orthogonal manner, we assessed the fluorescence of sfGFP under the control of P_Cym_/CymR, P_CymRC_/CymR_AM_ and P_CymRC_/CymR_GV_ circuits in response to increasing amounts of cumate. We observed that cumate circuits possessing identical CymR amino acid sequences, P_Cym_/CymR and P_CymRC_/CymR_GV_, resulted in similar kinetics within *B. cenocepacia* while P_CymRC_/CymR_AM_ showed heighten responsiveness to cumate even at low levels of induction (**Figure 3F**) with these trends also observed within *E. coli* (**Supplementary Figure 2**). Taken together, these results suggest that the use of P_CymRC_/CymR_AM_ leads to increased basal expression and narrower responsiveness to cumate compared to P_Cym_/CymR. These impacts are reversed by the restoration of amino acid substitutions within the CymR_AM_ sequence to that of the canonical form leading to reduced basal expression and widening of the cumate responsive range.

**Figure 3.**
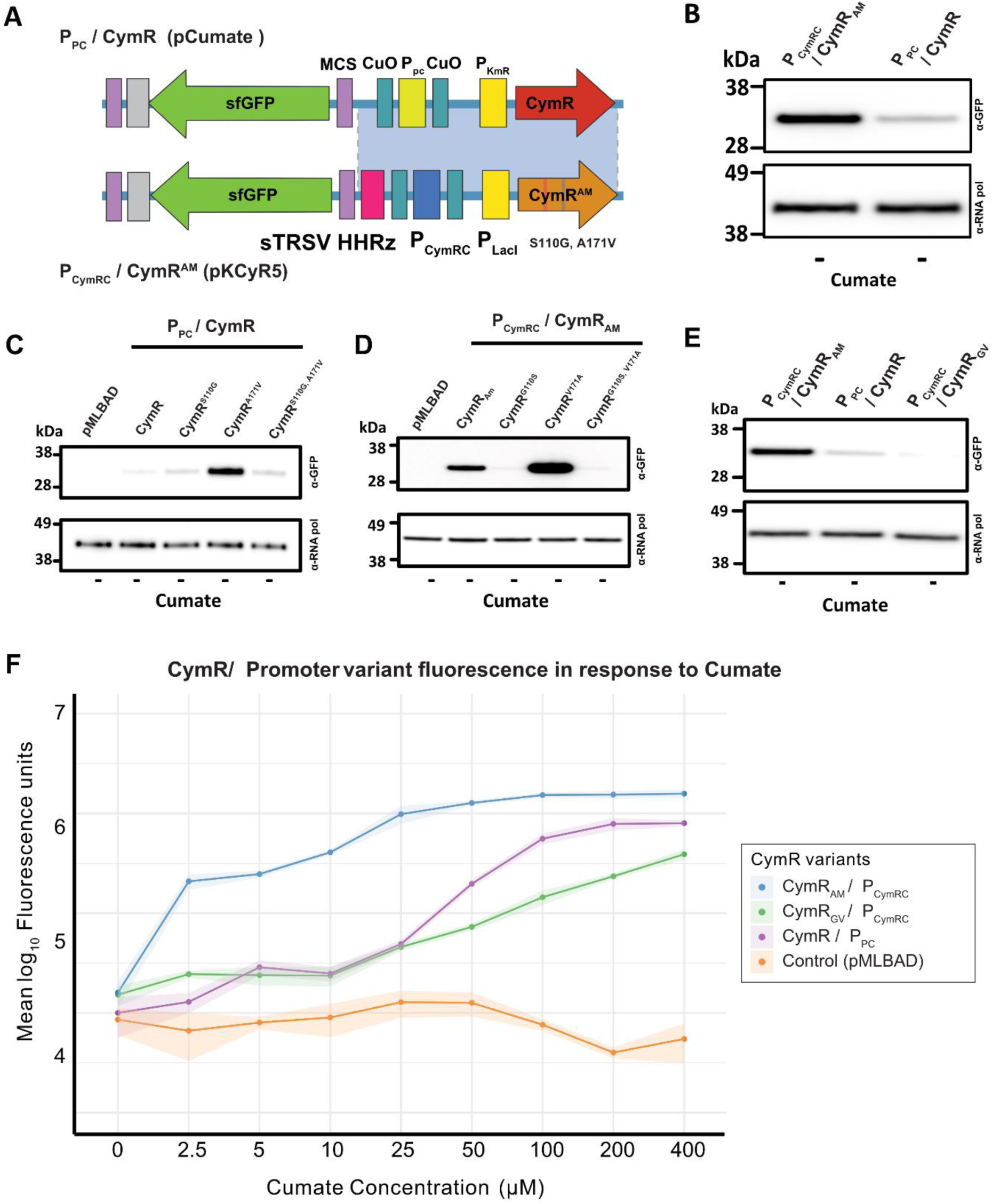
Assessment of basal expression across cumate circuits and the generation of the pCymRC/CymRGV cumate circuit. **A)** Schematic comparison between the regulatory circuits of generated cumate switch, P_pc_/CymR (top) and P_CymRC_/CymR_Am_ (bottom). **B)** Comparison of basal sfGFP expression between P_CymRC_/CymR_Am_ and the P_pc_/CymR variant. **C)** Western blot analysis of sfGFP expression using the wild-type repressor Ppc/CymR and its variants, revealing that the introduction of S110G or A171V mutations increases basal expression of sfGFP. **D)** Immunoblot analysis of CymR_Am_ derivatives, revealing that the reversion of G110S or V171A mutations reduces basal expression of sfGFP. **E)** sfGFP basal expression observed across the three cumate circuits, P_CymRc_/CymR_Am_, P_pc_/CymR, and p_CymRC_/CymR_GV_, reveals that p_CymRC_/CymR_GV_ demonstrates the lowest levels of basal expression. **F)** Line graph depicting sfGFP fluorescence intensity at varying cumate concentrations, with standard deviation represented by shaded areas (n = 3) of cumate circuits. All measurements were recorded 24 hours post-induction.

### P_CymRC_/CymR_GV_ allows enhanced tuneable control of protein expression and has minimal effects on the proteome of *B. cenocepacia*

While the reduced basal expression of sfGFP supports tighter regulatory control from P_CymRC_/CymR_GV_ compared to other cumate circuits, to directly assess this control within a biological context, we placed PglL under P_CymRC_/CymR_GV_ control, generating pCumate^GV^-PglL_Bc_- his, and again introduced this into *B. cenocepacia* Δ*pgl*L BCAL1086-his. Consistent with the reduced basal expression of PglL, western analysis revealed the absence of detectible BCAL1086-his without induction within strains containing pCumate^GV^-PglL_Bc_-his (**Figure 4A**). Upon cumate induction, we observe the appearance of BCAL1086-his at a similar mobility to BCAL1086-his observed with pCumate-PglL_Bc_-his, supporting the initiation of glycosylation (**Figure 4A**). Quantitative glycoproteomics/proteomics confirms the induction of glycosylation in response to cumate within *B. cenocepacia* Δ*pglL* BCAL1086-his containing pCumate^GV^-PglL_Bc_-his and pCumate-PglL_Bc_-his (**Figure 4B**). While the introduction of pCumate^GV^- PglL_Bc_-his reduces basal glycosylation, glycoproteomic analysis does support the presence of low-level glycosylation even in the absence of cumate induction (**Figure 4B, Supplementary Figure 3 & Supplementary Table 7**). While the presence of low-level glycosylation demonstrates that basal expression is still permissive from pCumate^GV^-PglL_Bc_-his, proteomic analysis reveals the lack of detectable PglL within strains containing pCumate^GV^-PglL_Bc_-his, compared to pCumate-PglL without induction (**Figure 4C, Supplementary Table 8**). Combined the absence of detectible BCAL1086-his and PglL within *B. cenocepacia* Δ*pglL* BCAL1086-his containing pCumate^GV^-PglL supports the reduced basal expression of PglL from P_CymRC_/CymR_GV_. By leveraging the improved control of P_CymRC_/CymR_GV_, we find this allows titratable control of glycosylation, as demonstrated by the inducible detection of BCAL1086-his in response to increasing cumate levels (**Figure 4D**). These results support that P_CymRC_/CymR_GV_ can be harnessed to allow precise tuneable protein expression within Burkholderia.

**Figure 4.**
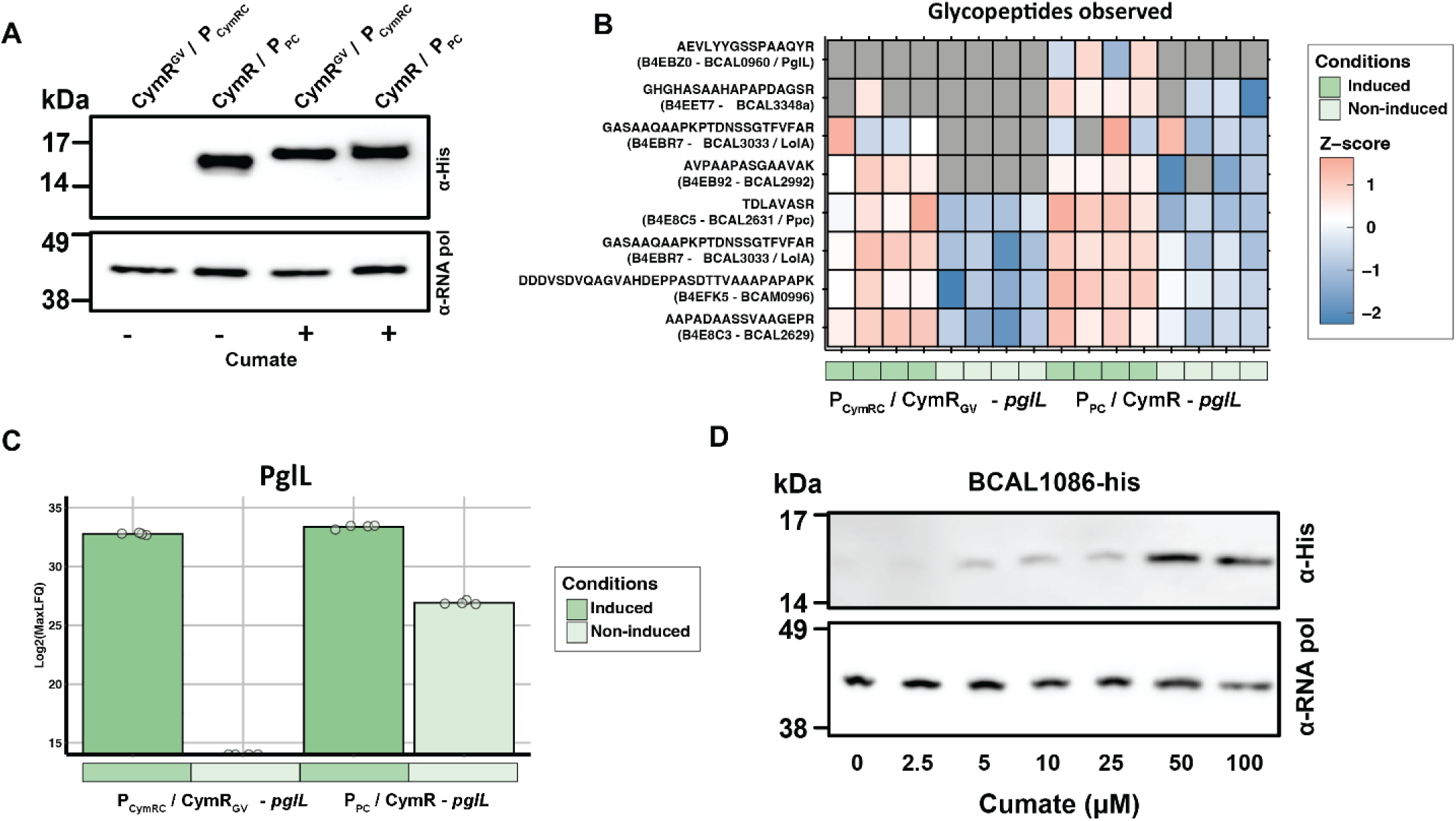
P_CymRC_/CymR_GV_ reduces basal expression, allowing inducible control of protein glycosylation. **A)** Immunoblot of BCAL1086-his supports improved control of protein glycosylation, with BCAL1086-his only detected upon induction of PglL-his under p_CymRC_/CymR_GV_ compared to P_pc_/CymR. **B)** Heatmap of selected glycopeptides observed within *B. cenocepacia* Δ*pgl*L BCAL1086-his, demonstrating reduced but detectable glycosylation from PglL under p_CymRC_/CymR_GV_ control compared to P_pc_/CymR (n = 4 per group). **C)** Quantification of PglL protein levels from whole-cell proteomic analysis reveals PglL is undetectable when under p_CymRC_/CymR_GV_ control in the absence of induction. **D)** Immunoblot of BCAL1086-his demonstrating the dose-dependent induction of BCAL1086-his expression with increasing cumate concentrations.

The ability to allow tight inducible control of protein expression makes P_CymRC_/CymR_GV_ a valuable tool, however, previous work within our lab has shown that inducers can drive large proteomic alterations within *B. cenocepacia* [43]. Thus, to understand the effects of cumate- based induction on the *B. cenocepacia* proteome, we undertook data-independent acquisition (DIA)-based proteomics comparing the impact of 100μM cumate on *B. cenocepacia K56-2* containing pCumate^GV^-sfGFP compared to *B. cenocepacia K56-2* containing the widely used rhamnose-inducible vector pScrhaB2 with 0.5% L-rhamnose [36]. A total of 4027 proteins were observed across groups with both cumate and rhamnose induction leading to notable changes in the proteome (**Figure 5A-B, Supplementary Table 9**). Interestingly, cumate induction resulted in reductions in the protein abundance of several proteins, associated with catabolic metabolism of aromatics (BCAM0810, BCAM0813, and BCAM0811), as well as reductions in known virulence factors including ZmpB (BCAM2307) and the Cable pilus-associated adhesin protein AdhA (BCAM2143) (**Figure 5A**). In contrast, rhamnose induction resulted in increased protein abundance of several proteins associated with stress-adaptation, including the putative exported BCAM2110 and Thio/disulfide interchange protein DsbC (pBCA043), as well as a decrease in the Alanine racemase DadX (BCAM1204), suggesting changes in metabolism in response to rhamnose induction (**Figure 5B**). Across these comparisons, we observed that overall cumate induction appears to lead to fewer proteomic alterations. Consistent with this, PCA analysis reveals tight clustering of pCumate^GV^-sfGFP groups yet demonstrates notable separation within the pScrhaB2 groups (**Figure 5C**). Finally, comparisons between the alterations observed in response to cumate and rhamnose induction reveal low overall overlap in the response to these inducers, supporting that cumate induction leads to discrete impacts on *B. cenocepacia* K56-2 (**Figure 5D**). Combined, these findings support that cumate induction drives minimal impacts on the *B. cenocepacia* K56-2 proteome and that the observed protein changes are orthogonal to the effects observed in response to rhamnose induction.

**Figure 5.**
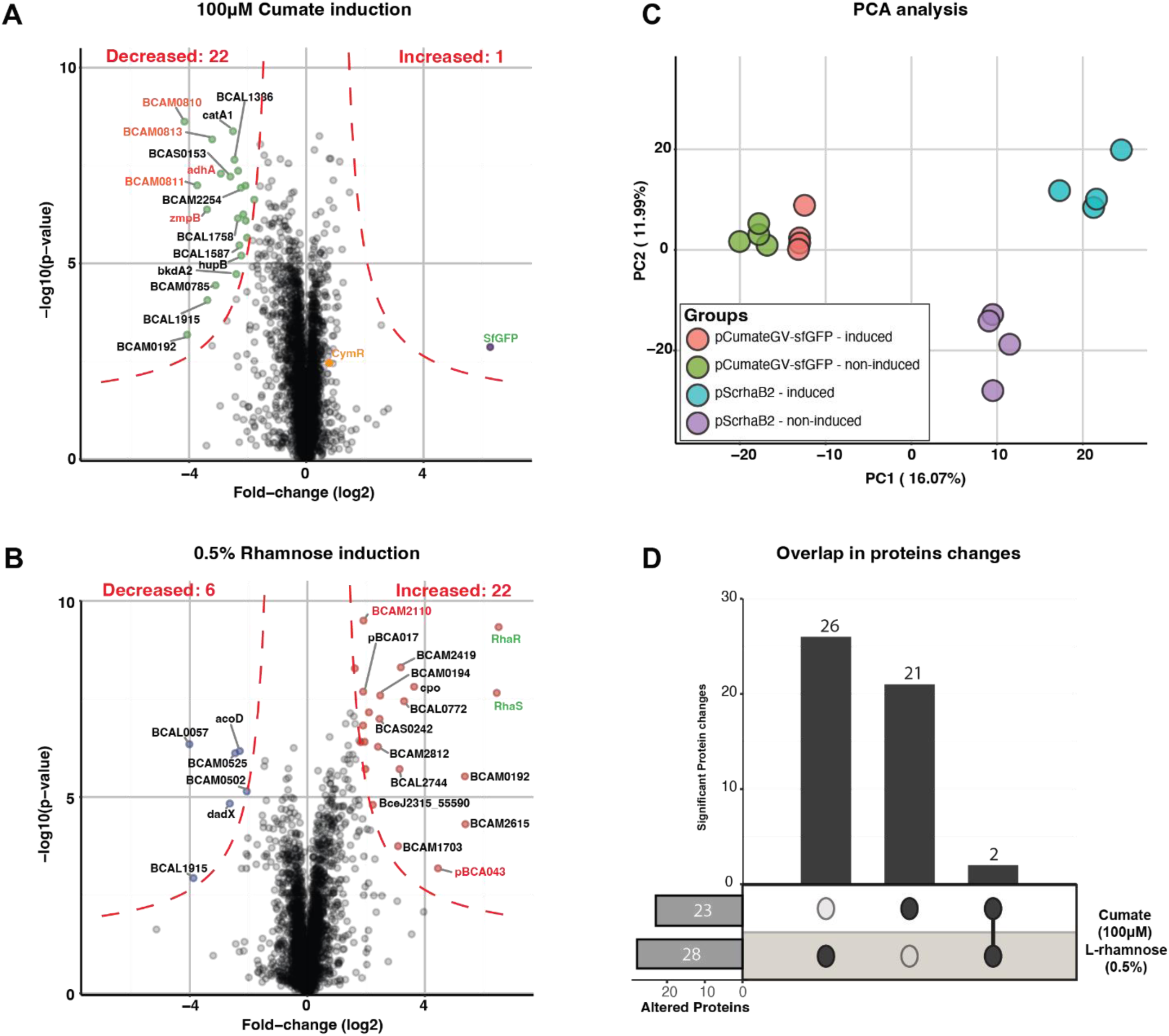
**Proteomic analysis of cumate and rhamnose induction within *B. cenocepacia*. A–B**) Volcano plots illustrating the differential protein abundance profiles between induced and non- induced conditions for cumate **A**) and rhamnose control **B**). Significance thresholds are denoted by the red S0 lines with altered proteins labelled. Proteins of interest are denoted in red, with the regulatory proteins RhaS/R denoted in green and CymR in orange. **C)** Principal component analysis (PCA) demonstrating distinct clustering of induced versus non-induced groups, confirming condition-dependent modest proteomic shifts. **D)** Upset plot showing the number of proteins exhibiting significant alterations under each condition, with overlap analysis highlighting shared and unique proteomic changes associated with cumate and rhamnose induction.

### Chromosomal P_CymRC_/CymR_GV_ allows inducible control within an *in vivo* context revealing the requirement of protein O-linked glycosylation for optimal intracellular replication

While the establishment of plasmid-based P_CymRC_/CymR_GV_ expression vector provides a versatile approach for protein induction, the requirement for continuous plasmid selection is not always ideal or achievable, such as within infection studies. To overcome these limitations, chromosomal integration provides an alternative strategy to allow tuneable expression under *in vivo* conditions. To assess the viability of using P_CymRC_/CymR_GV_ for *in vivo* chromosomal based protein expression, we introduced the P_CymRC_/CymR_GV_ circuit into a Mini-CTX1-based integration system [40, 46], leading to the generation of the vector pCTX-CymR^GV^ (**Figure 1C**). Using the Gentamicin-sensitive *B. cenocepacia K56-2* strain MH1K [47] we assessed intracellular burden within THP-1 cells after infection for 24-hours with OD_600_ normalised *B. cenocepacia* strains (**Supplementary Figure 4**) revealing no significant difference in bacterial colony forming units (CFUs) between MH1K CTX-sfGFP with or without cumate induction compared to MH1K (**Figure 6A**). Western analysis of infected THP-1 cells supports induction enables the controlled expression of sfGFP with sfGFP detectable as early as 2 hours post intracellular induction (**Figure 6B**). Utilising confocal immunofluorescence microscopy, we confirmed the internalisation of *B. cenocepacia* within THP-1 cells as well as the colocalization of sfGFP within *B. cenocepacia* in response to cumate induction post internalisation (**Figure 6C**). These observations support the ability of chromosomal integrated CTX P_CymRC_/CymR_GV_ to allow inducible expression of proteins within *B. cenocepacia* K56-2 during intracellular replication.

**Figure 6.**
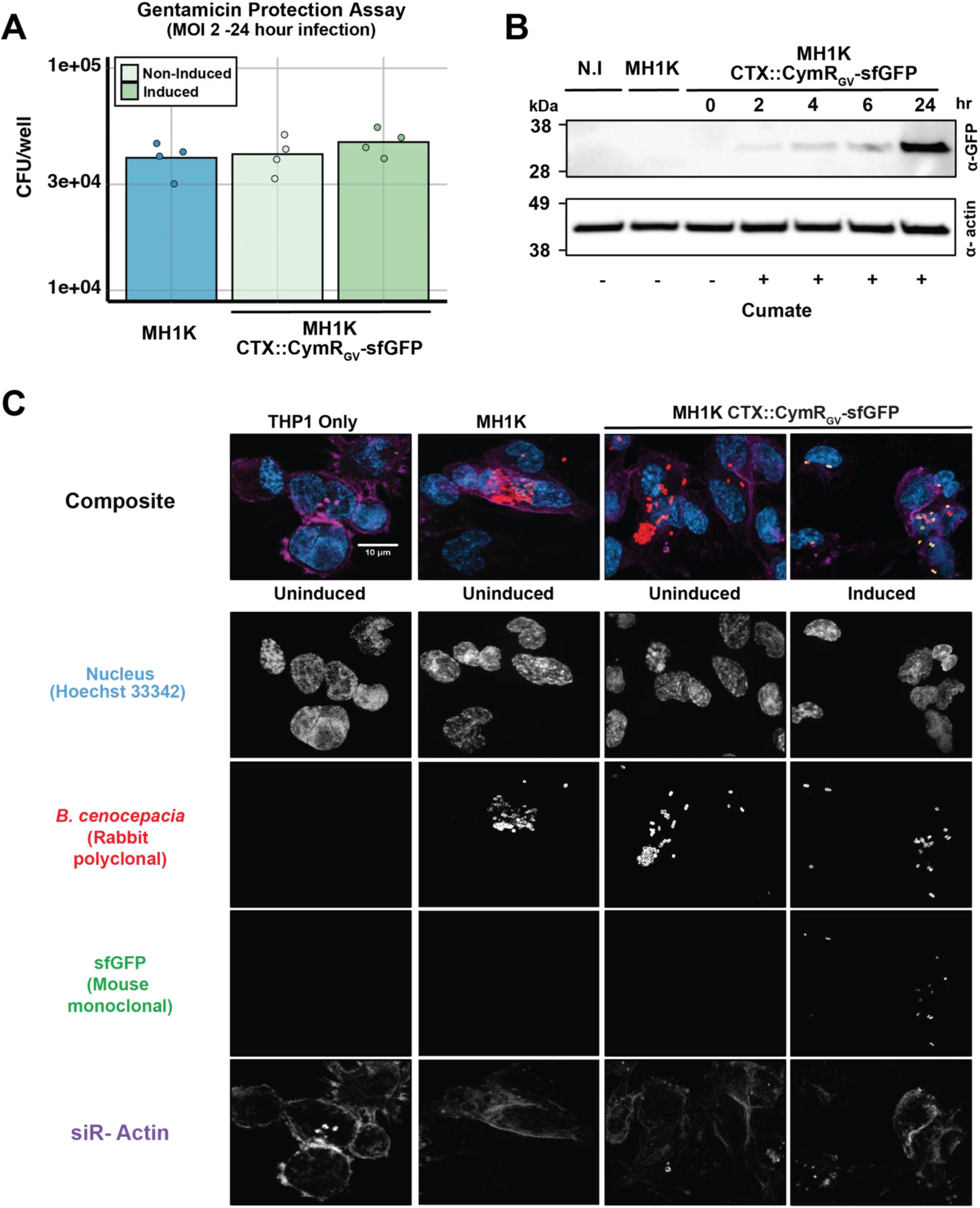
**Cumate induction allows inducible control of protein expression from *B. cenocepacia* within THP-1 cells**. A) Gentamicin protection assays demonstrate comparable bacterial burden, assessed as CFU/ml, within THP-1 cells under induced and non-induced conditions as compared to the MH1K strain. **B)** Immunoblot of sfGFP expression after 2, 4, 6, and 24 hours of cumate induction within THP-1 cells reveals detectible levels of sfGFP after 2 hours of induction. **C)** Immunofluorescence microscopy of internalised MH1K and MH1K containing CymR_GV_-sfGFP strains within THP-1 cells under induced and uninduced conditions reveal co-localisation of sfGFP and *B. cenocepacia* in an induction dependent manner. DNA has been stained with Hoechst 33342, *B. cenocepacia* with Alexa Fluor 568, sfGFP visualised through intrinsic fluorescence, and actin filaments labelled with SiR-actin 647.

Finally, to further demonstrate the capacity of pCTX-CymR^GV^ to facilitate intracellular studies of *B. cenocepacia*, we placed PglL*_Bc_*-his within pCTX-CymR^GV^ and integrated this into a Gentamicin-sensitive glycosylation-null strain, *B. cenocepacia* K56-2 Δ*amr*ABΔ*pglL*. Inducible expression of PglL*_Bc_*-his from chromosomally integrated CTX-CymR^GV^-PglL*_Bc_*-his was confirmed by western blotting (**Figure 7A**) and the successful restoration of glycosylation by glycoproteomic analysis (**Figure 7B**, **Supplementary Table 10 and 11**). Utilising Δ*amr*ABΔ*pglL* CTX-CymR^GV^-PglL*_Bc_*-his, we assessed the Intracellular burden of this strain within THP-1 cells 24-hour post infection compared to Δ*amr*AB and Δ*amr*ABΔ*pglL* using OD_600_ normalised infections (**Supplementary Figure 5**). CFU counts reveal a reduction in the bacterial load of Δ*pgl*L strains compared to Δ*amr*AB, with induction of PglL_Bc_-his within Δ*amr*ABΔ*pglL* CTX-CymR^GV^-PglL_Bc_-his restoring bacterial CFUs to parental levels (**Figure 7C**). Combined, these findings support that PglL*_Bc_*-his contributes to the ability of *B. cenocepacia* to replicate within THP-1 cells and demonstrates the ability of our CTX cumate-inducible system to facilitate inducible *in vivo* studies within *B. cenocepacia*.

**Figure 7.**
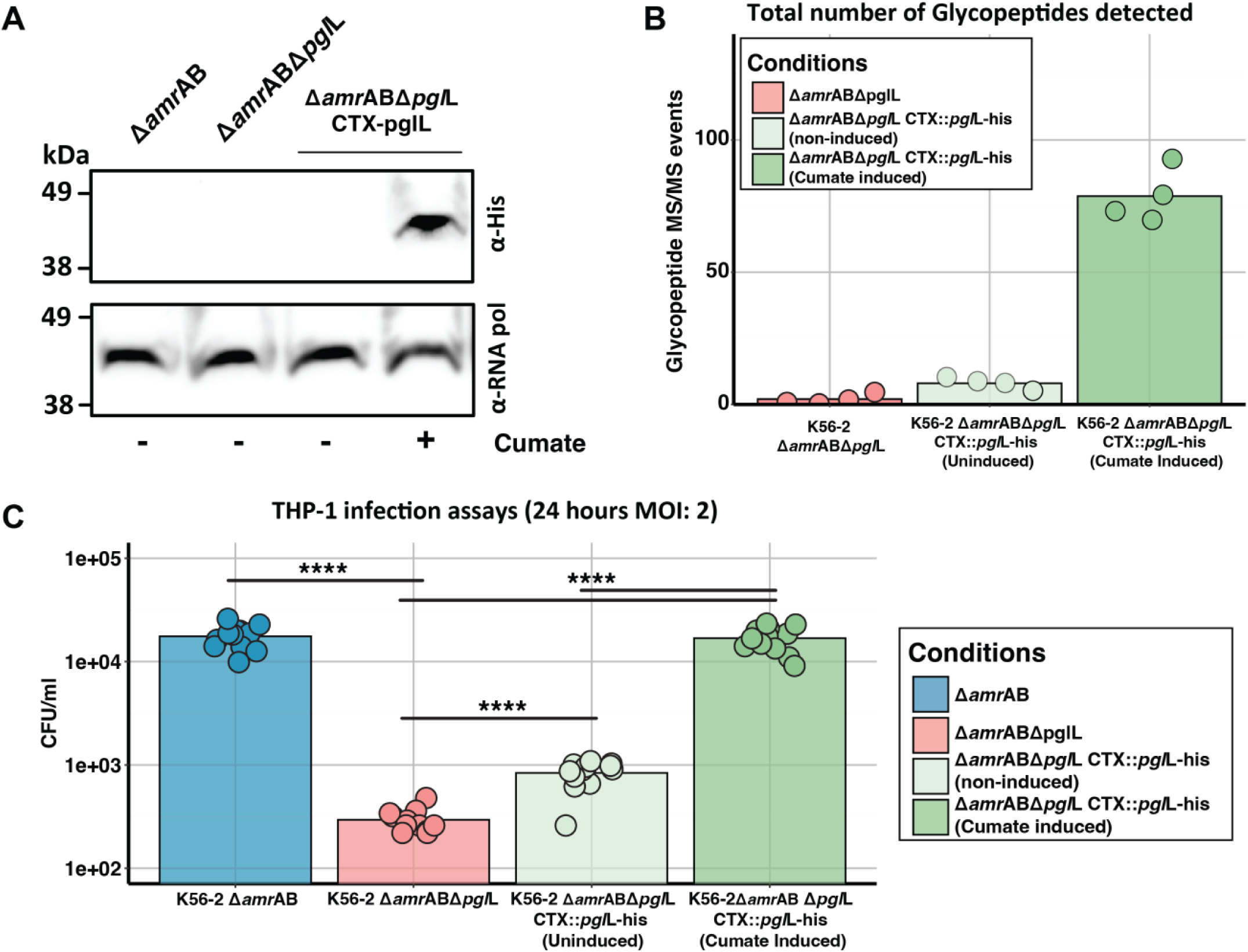
Protein glycosylation contributes to intracellular replication of *B. cenocepacia* within THP-1 cells. **A)** Immunoblot of His-tagged PglL in *B. cenocepacia* K56-2 Δ*amrAB*Δ*pglL* reveals induction dependent detection of PglL-his. **B)** Quantitation of glycopeptides identified by glycoproteomic analysis of *B. cenocepacia* K56-2 Δ*amr*ABΔ*pgl*L complemented with CymR_GV_- PglL_Bc_-his. Glycopeptides are undetectable in the Δ*amr*ABΔpglL mutant with cumate induction allowing the restoration glycosylation. **C)** Intracellular survival of *B. cenocepacia* K56-2 strains in THP-1 macrophages at 24 h post-infection (MOI = 2). The Δ*amr*ABΔ*pgl*L mutant exhibited reduced survival relative to Δ*amr*AB, while cumate induced pglL-his complementation restored intracellular burdent to near wild-type levels. (n=12 corresponding to 4 technical replicates of each of the three biological replicates undertaken, ****p<0.0001).

## Discussion

Inducible expression vectors enable controlled gene expression and are indispensable tools for molecular microbiology [2, 48]. While carbohydrate-inducible systems are widely employed within *B. cenocepacia*, these systems can be associated with several shortcomings, including modulating central metabolism and inducing stress responses [36, 42, 49]. To address these limitations, within this work, we have assessed and refined cumate-inducible systems for *B. cenocepacia* (**Figure 1**). While previous work demonstrated the functionality of the cumate- inducible system P_CymRC_/CymR_AM_ within *Burkholderia* [27], we find that the previously developed cumate circuits (P_Cym_/CymR & P_CymRC_/CymR_AM_) both possess high levels of basal expression, limiting their utility and motivating us to improve the available circuits which are utilised for cumate induction. Critically, we find that while the functionality of cumate circuits can be assessed using reporter proteins by western or fluorescent assays (**Figure 2 & 3**), these approaches are largely insufficient to gauge basal expression. As has been exploited to allow highly sensitive detection of protein translocation [50, 51], we observed that the use of an enzymatic process, in this case protein glycosylation, provides a more sensitive and meaningful way to track basal expression within *B. cenocepacia,* allowing us to refine a new cumate circuit (P_CymRC_/CymR_GV_) utilising western-based analysis with the glycosylation-sensitive protein BCAL1086-his [43, 45] and glycoproteomics.

Our observations of high basal expression utilizing the P_Cym_/CymR circuit are in line with prior reports noting challenges with this inducible system when manipulating deleterious enzymatic processes [52, 53]. Pöschel *et al* noted that within *Methylorubrum extorquens*, the use of P_Cym_/CymR for the expression of a recombinant mevalonate pathway resulted in growth abnormalities, which were only suppressed by a spontaneous mutation within the promoter P_Cym_, reducing basal expression [53]. Our work supports that while P_Cym_/CymR is functional in *B. cenocepacia*, it does not provide sufficient control to limit basal expression (**Figure 2**). While reducing basal expression motivated us to explore the use of P_CymRC_/CymR_AM_, which was developed to possess improved characteristics [3], we find that this circuit, both within *B. cenocepacia* (**Figure 3**) and *E. coli* **(Supplementary Figure 1 & 2),** displays shallower cumate responsiveness and higher levels of basal expression compared to the canonical P_Cym_/CymR circuit. These characteristics can be reverted by restoring CymR_AM_ to the progenitor CymR sequence (**Figure 3 & 4**), highlighting an effective means to improve basal control within vectors that have utilized P_CymRC_/CymR_AM_. Utilizing the P_CymRC_/CymR_GV_ circuit, we find that tight expression control can be achieved from both plasmid (**Figure 4**) and chromosomally integrated (**Figure 6 & 7**) vectors within *B. cenocepacia*. While our work has predominantly focused on *B. cenocepacia*, these systems are equally amenable to other members of the *Burkholderia* genus, including *Burkholderia thailandensis*, where we have confirmed the functionality of the P_CymRC_/CymR_GV_ circuit (**Supplementary Figure 6**). Importantly, it should be noted that while many *Burkholderia* species lack pathways for the catabolic metabolism of cumate, several species do possess cym pathways which have been shown to be functional, such as *Burkholderia xenovarans* [54, 55]. Thus, cumate induction and responsiveness should be assessed on a case-by-case basis when applying these systems to diverse *Burkholderia* members as the assumption that cumate is non-metabolizable may not hold true for all species.

A key motivation for refining cumate-inducible systems for use in *Burkholderia* was driven by our observation of the proteomic consequences of rhamnose induction [43]. Consistent with our prior reports, rhamnose induction drives both increases and decreases in the *B. cenocepacia* proteome (**Figure 5B**). While we observe that cumate induction also results in notable effects on the proteome, these effects are discrete from those observed with rhamnose (**Figure 5D**) and the magnitude of these effects appear less extreme than those observed with rhamnose (**Figure 5A & C**). Interestingly, many observed protein changes upon cumate induction appear to be associated with a reduction in protein levels, including reductions in known virulence factors such as ZmpB/BCAM2307 and the Cable pilus-associated adhesin protein adhA/BCAM2143. While this suggests that the addition of cumate may suppress potential virulence pathways within *B. cenocepacia*, this did not lead to alterations in intracellular survival within strains containing chromosomally integrated P_CymRC_/CymR_GV_ vectors (**Figure 6A & 7C**). Critically, as cumate is cell-permeable, we demonstrate that this provides the unique ability to trigger protein induction during *in vivo* studies (**Figure 6C**). While approaches for intracellular expression have been developed for other bacterial pathogens, such as *Salmonella*, where promoters associated with the *Salmonella* pathogenicity island 2 have been used for temporal protein control [56, 57], these systems are not generalizable across bacterial species. To our knowledge, this is the first example of using cumate-induced systems for titratable protein expression control within an intracellular bacterial infection model. Thus, while cumate induction has been used for eukaryotic expression systems [26], its ability to allow expression induction within an infection context expands the field’s capacity to study intracellular pathogens such as *B. cenocepacia*.

Finally, we demonstrate that the loss of glycosylation reduces the bacterial burden within a THP-1 infection model, and utilizing our refined cumate-inducible system, this effect can be reversed upon complementation (**Figure 7C**). This finding is consistent with a growing body of evidence that *Burkholderia* O-glycosylation contributes to fitness and pathogenicity [58, 59]. Interestingly, while protein glycosylation has been shown by several teams to contribute to *B. cenocepacia* survival in *Galleria mellonella* [60, 61], the observation of its role in mammalian intracellular replication has not been previously demonstrated. Recently, Paszti *et al* demonstrated using Tn-Seq that several genes associated with protein glycosylation glycan biosynthesis (BCAL3114 & BCAL3115) were essential within both *G. mellonella* and *ex vivo* pig lung models; however, they only observed the requirement for PglL_Bc_ (BCAL0960) within *G. mellonella* [60]. Our observation that the lack of PglL_Bc_ reduces but does not abolish the recovery of *B. cenocepacia* from THP-1 suggests that O-linked glycosylation contributes to survival, but may not be essential for pathogenesis per se. Utilizing this inducible system, future work will seek to understand how the absence of glycosylation shapes the course of intracellular infections and if the lack of glycosylation simply reduces bacterial replication or impacts specific processes during intracellular survival, leading to the observed reduction in bacterial load.

Combined, this work establishes a set of refined cumate-inducible vectors for *B. cenocepacia*, allowing tight inducible control as well as the capacity to induce protein expression within *in vivo* models. While the focus of this work sought to expand the availability of inducible systems for *Burkholderia* studies, these cumate vectors are likely applicable to other non-model bacteria where gene control can be challenging. In summary, by refining and developing novel cumate- inducible vectors, this work expands the genetic toolkit for studying the *Burkholderia* genus.

## Material and Methods

### Bacterial strains, Plasmids and growth conditions

The bacterial strains utilised in this study are detailed in **Supplementary Table 1**. The Plasmids used in this study are listed in **Supplementary Table 2**. Strains were cultured in Lysogeny Broth (LB- BD™, USA) or on LB agar (1.5–2% w/v), prepared according to the manufacturer’s instructions and containing 0.5% NaCl. Liquid cultures were incubated overnight at 37 °C with shaking, whereas agar plates were incubated at 37 °C overnight for *E. coli* and for 24–72 hours for *B. cenocepacia*. Antibiotics were incorporated into cultures to select plasmids/transconjugants at a final concentration of 50 μg/mL for *E. coli*, 100 μg/mL for *B. cenocepacia for* trimethoprim; 20 μg/mL for *E. co*li, 150 μg/mL for *B. cenocepacia for* tetracycline as well as 50 μg/mL kanamycin and 10 μg/mL gentamicin for *E. coli*. To selectively inhibit helper and donor *E. coli* strains after triparental mating, ampicillin was added to agar plates at a concentration of 100 μg/mL and polymyxin B at 25 µg/mL [34]. Culturing of strains harbouring the temperature-sensitive plasmid pFlp-Ab5 was performed at 30 °C, followed by a shift to 37 °C to facilitate plasmid curing [62]. Induction of *B. cenocepacia* strains was conducted by adding filter-sterilised L-rhamnose monohydrate (Sigma) to LB broth at 0.5% rhamnose or cumate (4-isopropylbenzoic acid, Sigma) dissolved in 100% ethanol and added at concentrations ranging from 2.5 μM to 400 μM. An equivalent volume of solvent (sterile water for rhamnose and ethanol for cumate) was added to the culture media for non-induced controls.

### Recombinant DNA methods

The oligonucleotides utilized in this study are detailed in **Supplementary Table 3**. All PCR amplifications for cloning were performed using Q5 DNA polymerase (New England Biolabs, USA), with the inclusion of 2% DMSO for the amplification of *B. cenocepacia* DNA owing to its high GC content. Genomic DNA was isolated using the EZ-10 Spin Column Bacterial Genomic DNA Mini-Preps Kit (Bio Basic, Canada). PCR cleanup and restriction digest purifications were conducted with the Zymoclean Gel DNA Recovery Kit (ZymoResearch, USA). Plasmid DNA was isolated using the QIAprep Spin Miniprep Kit (Qiagen, Germany). Vectors were generated by amplifying regions of interest and then assembling amplicons with the aid of Gibson Assembly [63] using the NEBuilder® HiFi DNA Assembly Mix (New England Biolabs). Before Gibson Assembly DNA fragments were evaluated for correctness based on size through agarose gel electrophoresis. Single nucleotide mutations were generated using Q5 PCR-based site-directed mutagenesis, followed by DpnI (New England Biolabs) treatment to remove template DNA. Plasmid mixtures were introduced into chemically competent *E. coli* PIR2 cells through heat shock transformation. The cells were plated on selective media and screened via colony PCR using GoTaq® Green Master Mix (Promega, USA). All plasmids utilized in this study were verified by either Sanger sequencing using the Australian Genome Research Facility (Melbourne, Australia) or nanopore plasmid sequencing performed by Plasmidsaurus (SNPsaurus LLC, USA). A summary of all plasmids and the primer combinations used for their construction are provided in **Supplementary Table 2**.

### Conjugation of plasmids into *B. cenocepacia*

Plasmids were introduced into *B. cenocepacia* K56–2 strains utilising triparental mating [34] and *E. coli* PIR2 strains carrying donor plasmids. Conjugations were allowed to proceed for 24 hours, then successful transconjugants were selected with trimethoprim or tetracycline, and the introduction of plasmids of interest confirmed using PCR-based screening. For the generation of strains containing CTX integration vectors [40] the tetracycline resistance marker was excised by the introduction of the temperature sensitive flippase vector pFLP-Ab5 [34] with excision confirmed using antibiotic screening and PCR based confirmation. pFLP-Ab5 plasmid curing was undertaken without selection at 37 °C.

### Growth conditions and induction assays

sfGFP fluorescence was quantified in *E. coli* and *B. cenocepacia* utilising a CLARIOstar® Plus multimode plate reader (BMG Labtech, Germany) with black, flat-bottom 96-well plates (200 µL per well, n = 3). Cultures were inoculated with varying concentrations of cumate (0–200 µM for *E. coli*, 0–400 µM for *B. cenocepacia*) and incubated at 37 °C. Optical density was measured at 600 nm to assess cell density, and GFP fluorescence was quantified with an excitation at 485 nm and emission between 520 and 530 nm. Measurements were taken at 10-minute intervals over a 24-hour period, with orbital shaking applied before each measurement to ensure uniformity. Background signals from controls were subtracted using blank values.

### Western Blotting

Overnight bacterial cultures (1ml of stationary phase, OD_600nm_ ∼1.5) were pelleted then boiled in 1X SDS-PAGE sample buffer (2% SDS, 10% glycerol, 5% 2- mercaptoethanol, 0.004% bromphenol blue and 75 mM Tris HCl, pH 6.8) for 10 minutes. Protein lysates were separated on NuPAGE 4-12% Bis-Tris gradient polyacrylamide gels in MES buffer (Thermo Fisher Scientific). Proteins were transferred to a nitrocellulose membrane using the iBlot 3 Western transfer system (Thermo Fisher Scientific) and blocked for 1 hr in 5% skim milk in TBS-T (Tris buffered saline with 0.1% Tween-20). Nitrocellulose membranes were probed with either mouse α-histidine tag (1:1000; clone: AD1.1.10, Bio-Rad) or mouse α-GFP (1:5000; IgG1κ clones 7.1 and 13.1, 11814460001, Sigma) overnight at 4 °C, washed three times with TBS-T, then incubated with horseradish peroxidase (HRP) conjugated α-mouse antibody (1:5000; NEF82200-1EA, PerkinElmer) for 1 hr at room temperature. Proteins were detected using Clarity Western ECL Substrate (Bio-Rad) and images obtained using an Amersham imager 800 (GE Healthcare Life Sciences). Membranes were Stripped with Restore™ Plus Stripping Buffer for 10 minutes before being re-blocked in 5% skim milk in TBS-T and probed with mouse α-*E. coli* RNA polymerase primary antibody (1:5000; clone: 4RA2, Neoclone) or mouse α-Actin (1:5000; clone: BA3R, Thermo Fisher Scientific) followed by HRP conjugated α-mouse and imaged as above. All antibodies were diluted in TBS-T with 1% bovine serum albumin (BSA; Sigma-Aldrich).

### Gentamicin protection assays

Gentamicin protection assays were undertaken using the approach of Hamad *et al* [47]. Briefly, THP-1 cells (ATCC TIB-202) grown in RPMI-1640 culture media (Gibco, Thermo Fisher) supplemented with 10% Fetal Bovine Serum (FBS) at 37 °C with 5% CO_2_ were seeded in twenty-four-well plates at a final concentration of 3.0×10^5^ and differentiated with 50 ng/mL Phorbol 12-myristate 13-acetate for 72 hours prior to use in infection assays. Overnight cultures of *B. cenocepacia* were standardised to an OD_600_ of 1.0 (∼1x10^9^ cells per mL) and washed three times with 1 mL of pre-warmed RPMI-1640 with 10% FBS. Bacterial cells were resuspended into 1 mL of fresh RPMI + 10% FBS and 100µl of 6.0×10^5^ cfu/ml, corresponding to a multiplicity of Infection of 2, then added to the differentiated THP-1 cells. To synchronise and enhance bacterial uptake plates were centrifuged for 1 min at 300*g* and incubated for 1 hour at 37 °C under 5% CO_2_. Infected macrophages were washed three times with PBS to remove external bacteria. The media was then replaced with fresh RPMI + 10% FBS containing 50 μg/mL gentamicin for 30 minutes, then the media replaced with RPMI + 10% FBS containing 10 μg/mL gentamicin. For in vivo induction, infected THP 1 cells were maintained in RPMI media containing 10% FBS,10 μg/mL gentamicin and 100μM cumate following 30 minutes of 50 μg/mL gentamicin killing. After a total of 24hours from the initial addition of bacteria, cells were washed once with PBS then lysed with 1% Triton X-100 in PBS. Recovered intracellular bacteria were enumerated by plating serial dilutions on LB agar plates.

### Immunofluorescence microscopy

THP-1 cells were seeded then differentiated on coverslips in 24-well plates as above prior to infection with *B. cenocepacia* strains at an MOI of 2. 24 hours post-infection, the culture media was removed and cells washed with PBS then fixed by the addition of 4% pre-chilled paraformaldehyde and incubated at room temperature for 15 minutes. Fixed cells were washed with PBS five times then permeabilized in PBS containing 1% BSA and 0.2% Triton X-100 for 1 hour, before blocking in PBS containing 1% BSA and 0.1% Tween-20 for 1 hour. Cells were stained with α-*Burkholderia* polyclonal rabbit serum (1:500 in PBS with 1% BSA- generated from lysates of *B. cenocepacia K56-2* through the Walter and Eliza Hall Institute antibody facility) for 1 hour. Unbound antibodies were removed by washing with PBS containing 1% BSA and 0.1% Tween-20 three times for 10 minutes and coverslips then incubated with α-rabbit Alexa Fluor 568 secondary antibody (1:2000 in PBS containing 1% BSA and 0.1% Tween-20) for 1 hour in the dark. Unbound secondary antibody was removed by washing three times with PBS containing 1% BSA and 0.1% Tween-20 three times for 10 minutes before the addition of DNA and actin stains Hoechst 33342 (0.1 µg/mL, Abcam) and SiR-Actin (100 nM, Spirochrome) for 20 minutes in PBS containing 1% BSA. Coverslips were then washed five times with PBS containing 1% BSA and 0.1% Tween-20 before being mounted using ProLong™ Gold Antifade (Invitrogen) on a glass slide and allowed to cure overnight and subsequently sealed with nail polish. Imaging was conducted utilizing an LSM980 confocal microscope (Zeiss, Germany) using the smart setup acquisition mode and an oil immersion 63X objective. Z-stack images were reconstructed through orthogonal projection, and visualization was performed using Fiji software [64] and three independent infection assays undertaken. Image processing was restricted to linear modifications of brightness and contrast, uniformly applied to all samples.

### Proteomic sample preparation

*B. cenocepacia* cultures for proteomic analysis were incubated overnight with or without inducers (Cumate or Rhamnose), under shaking conditions at 180 rpm. Cultures were adjusted to an OD₆₀₀ of 1.0, harvested by centrifugation at 10,000 × g for 10 minutes at 4 °C, washed three times with ice-cold PBS, and snap-frozen at −80 °C until further processing. Frozen whole-cell samples were solubilised in 4% SDS, 100mM Tris pH 8.5 by boiling for 10 min at 95 °C then protein concentrations assessed by bicinchoninic acid protein assays (Thermo Fisher Scientific). 100μg of each biological replicate/ sample was prepared for digestion using S-traps mini columns (Protifi, USA) according to the manufacturer’s instructions. Briefly, samples were reduced with 10mM Dithiothreitol for 10 mins at 95 °C and then alkylated with 50mM Iodoacetamide in the dark for 1 hour. Samples were acidified to 1.2% phosphoric acid and diluted with seven volumes of S-trap wash buffer (90% methanol, 100mM Tetraethylammonium bromide pH 7.1) before being loaded onto S-traps and washed 3 times with S-trap wash buffer. Samples were then digested with 2 μg of SoluTrypsin (Sigma) overnight at 37 °C before being collected by centrifugation with washes of 100mM Tetraethylammonium bromide, followed by 0.2% formic acid followed by 0.2% formic acid / 50% acetonitrile. Samples were dried down and further cleaned up using C_18_ Stage [65, 66] tips to ensure the removal of any particulate matter.

### Data-Dependent Acquisition (DDA) LC-MS analysis

Combined Glycopeptide and proteomic analysis were undertaken using DDA based analysis. Samples were re-suspended in Buffer A* (2% acetonitrile, 0.1% trifluoroacetic acid) and separated using a two-column chromatography setup composed of a PepMap100 C_18_ 20-mm by 75-μm trap (Thermo Fisher Scientific) and a PepMap C_18_ 500-mm by 75-μm analytical column (Thermo Fisher Scientific) using a Dionex Ultimate 3000 UPLC (Thermo Fisher Scientific). Samples were concentrated onto the trap column at 5 μl/min for 6 min with Buffer A (0.1% formic acid, 2% DMSO) and then infused into an Orbitrap Fusion Lumos™ (Thermo Fisher Scientific) at 300 nl/minute via the analytical column. Peptides were separated by altering the buffer composition from 3% Buffer B (0.1% formic acid, 77.9% acetonitrile, 2% DMSO) to 28% B over 85 minutes, then from 23% B to 40% B over 4 minutes and then from 40% B to 80% B over 3 minutes. The composition was held at 80% B for 2 minutes before being returned to 3% B for 10 minutes. The Orbitrap Lumos Mass Spectrometer was operated in a data-dependent mode automatically switching between the acquisition of a single Orbitrap MS scan (300-2000 m/z, maximal injection time of 25 ms, an Automated Gain Control (AGC) set to a maximum of 300% and a resolution of 60k) and 3 seconds of Orbitrap MS/MS HCD scans of precursors (NCE of 35%, a maximal injection time of 54 ms, a AGC of 100% and a resolution of 15). HCD scans containing HexNAc-associated oxonium ions (204.0867, 138.0545 and 366.1396 m/z) triggered two additional product-dependent MS/MS scans [67] of potential glycopeptides; an Orbitrap EThcD scan (NCE 15%, maximal injection time of 150 ms, AGC set to a maximum of 250% ions with a resolution of 30k using the extended mass range setting to improve the detection of high mass glycopeptide fragment ions [68]) and a stepped collision energy HCD scan (using NCE 35% with 5% Stepping, maximal injection time of 150 ms, an AGC set to a maximum of 250% ions and a resolution of 30k).

### Data-Independent Acquisition (DIA) LC-MS analysis

Total proteome analysis was undertaken using DIA based analysis. C_18_ enriched proteome samples were re-suspended in Buffer A* and separated using a two-column chromatography setup composed of a PepMap100 C_18_ 20-mm by 75-μm trap (Thermo Fisher Scientific) and a PepMap C_18_ 500-mm by 75-μm analytical column (Thermo Fisher Scientific) using a Dionex Ultimate 3000 UPLC (Thermo Fisher Scientific). Samples were concentrated onto the trap column at 5 μl/min for 6 min with Buffer A (0.1% formic acid, 2% DMSO) and then infused into an Orbitrap Eclipse™ (Thermo Fisher Scientific) at 300 nl/minute via the analytical column. Peptides were separated by altering the buffer composition from 3% Buffer B (0.1% formic acid, 77.9% acetonitrile, 2% DMSO) to 28% B over 70 minutes, then from 23% B to 40% B over 4 minutes and then from 40% B to 80% B over 3 minutes. The composition was held at 80% B for 2 minutes before being returned to 3% B for 10 minutes. The Orbitrap Eclipse was operated in a data-independent mode automatically switching between the acquisition of a single Orbitrap MS1 event (120k resolution, AGC set to a maximum of 100%, 350-1400 m/z) and 50 MS2 scans (NCE 30%, 30k resolution, AGC set to a maximum of 1000%, 200-2000 m/z and a maximal injection time of 55 ms) of a width of 13.7 m/z collected over the mass range of 361 to 1033 m/z.

### DDA-based proteomic analysis

DDA datasets were analysed using MSFragger (version 22.0) [69–71]. Samples were searched with a Tryptic specificity, allowing a maximum of two missed cleavage events and Carbamidomethyl set as a fixed modification of Cysteine while oxidation of Methionine allowed as a variable modification. The *Burkholderia* glycans HexHexNAc_2_ (elemental composition: C_22_O_15_H_36_N_2_, mass: 568.2115 Da) and Suc-HexHexNAc_2_ (elemental composition: C_26_O_18_H_40_N_2_, mass: 668.2276 Da) were included as variable modifications at Serine in line with the strong preference for PglL glycosylation at Serine residues [58, 59]. The glycan fragment ions were defined as 204.0866, 186.0760 168.0655, 366.1395, 144.0656, 138.0550, 466.1555 and 407.1594. A maximum mass precursor tolerance of 20 ppm was allowed at both the MS1 and MS2 levels. Samples were searched against the *B. cenocepacia* reference proteome J2315 (Uniprot accession: UP000001035, 6,993 proteins) [72] supplemented with the proteins RhaS (HTH-type transcriptional activator RhaS, Uniprot accession: P09377), RhaR (HTH-type transcriptional activator RhaR, Uniprot accession: P09378), Uniprot accession: CymR, Uniprot accession: O33453). MS1-based quantitation was undertaken using MaxLFQ [73] with matching between runs enabled. Analysis of protein abundances was undertaken using Perseus [74] with data visualization was undertaken using ggplot2 [75] in R.

### DIA-based proteomic analysis

DIA data was searched with MSFragger (version 22.0) [69–71] using the “DIA_SpecLib_Quant” workflow utilizing DIA-NN [76]. Samples were searched with a Tryptic specificity, allowing a maximum of two missed cleavage events and Carbamidomethyl set as a fixed modification of Cysteine while oxidation of Methionine allowed as a variable modification. A maximum mass precursor tolerance of 20 ppm was allowed at both the MS1 and MS2 levels. Samples were searched against the *B. cenocepacia* reference proteome J2315 (Uniprot accession: UP000001035, 6,993 proteins) [72] supplemented with the proteins RhaS (HTH-type transcriptional activator RhaS, Uniprot accession: P09377), RhaR (HTH-type transcriptional activator RhaR, Uniprot accession: P09378), Uniprot accession: Uniprot accession: O33453). DIA-NN-based quantitation was used for statistical analysis of protein abundances and analysed within Perseus [74]. Missing values were imputed based on the total observed protein intensities with a range of 0.3 σ and a downshift of 2.5 σ with biological replicates grouped together to allow comparison by student t-tests. Proteins were considered altered if they satisfied both p-value and fold change alterations as defined by the function y = c/(x - x0) where c = curvature and x0 = minimum fold change with were x0 = 1 and c = 4 [77]. Data visualization was undertaken using ggplot2 [75] in R.

### Data Availability

All raw mass spectrometry data, analysis outputs, and scripts have been deposited in the PRIDE ProteomeXchange repository. Accession numbers and experiment details are provided in Supplementary Table 4.

## Acknowledgements

N.E.S was supported by an Australian Research Council (ARC) Future Fellowship (FT200100270), an ARC Discovery Project Grant (DP210100362) and a National Health and Medical Research Council Ideas grant (2018980). We thank the Melbourne Mass Spectrometry and Proteomics Facility of The Bio21 Molecular Science and Biotechnology Institute for access to MS instrumentation. We gratefully acknowledge the Biological Optical Microscopy Platform (https://microscopy.unimelb.edu.au/om) at the University of Melbourne, for their support and assistance with microscopy. We thank Nicholas Coleman for sharing the plasmid pUS250-sfGFP and Tung Hoang for sharing the plasmid pFlp-Ab5.

## References

1. Alper, H., et al., Tuning genetic control through promoter engineering. Proceedings of the National Academy of Sciences, 2005. 102(36): p. 12678–12683.

2. Khalil, A.S. and J.J. Collins, Synthetic biology: applications come of age. Nature Reviews Genetics, 2010. 11(5): p. 367–379.

3. Meyer, A.J., et al., Escherichia coli "Marionette" strains with 12 highly optimized small-molecule sensors. Nat Chem Biol, 2019. 15(2): p. 196–204.

4. Cardona S.T., Carmen, M.L. and Valvano M. A., Identification of Essential Operons with a Rhamnose-Inducible Promoter in Burkholderia cenocepacia. Applied and Environmental Microbiology, 2006. 72(4): p. 2547–2555.

5. Chen, Y., et al., Tuning the dynamic range of bacterial promoters regulated by ligand-inducible transcription factors. Nature Communications, 2018. 9(1): p. 64.

6. Kent, R. and N. Dixon, Contemporary Tools for Regulating Gene Expression in Bacteria. Trends Biotechnol, 2020. 38(3): p. 316–333.

7. Lutz, R. and H. Bujard, Independent and tight regulation of transcriptional units in Escherichia coli via the LacR/O, the TetR/O and AraC/I1-I2 regulatory elements. Nucleic acids research, 1997. 25(6): p. 1203–1210.

8. Guzman, L.-M., et al., Tight regulation, modulation, and high-level expression by vectors containing the arabinose PBAD promoter. Journal of bacteriology, 1995. 177(14): p. 4121–4130.

9. Carrillo Rincón, A.F., et al., A dual-inducible control system for multistep biosynthetic pathways. Journal of Biological Engineering, 2024. 18(1): p. 68.

10. Yu, T.C., et al., Multiplexed characterization of rationally designed promoter architectures deconstructs combinatorial logic for IPTG-inducible systems. Nature Communications, 2021. 12(1): p. 325.

11. Fuller, F., A family of cloning vectors containing the lacUV5 promoter. Gene, 1982. 19(1): p. 43–54.

12. Skerra, A., Use of the tetracycline promoter for the tightly regulated production of a murine antibody fragment in Escherichia coli. Gene, 1994. 151(1-2): p. 131–5.

13. Haldimann, A., L.L. Daniels, and B.L. Wanner, Use of new methods for construction of tightly regulated arabinose and rhamnose promoter fusions in studies of the Escherichia coli phosphate regulon. J Bacteriol, 1998. 180(5): p. 1277–86.

14. Hothersall, J., et al., The PAR promoter expression system: Modified lac promoters for controlled recombinant protein production in Escherichia coli. N Biotechnol, 2021. 64: p. 1–8.

15. Li, Y., et al., A cross-species inducible system for enhanced protein expression and multiplexed metabolic pathway fine-tuning in bacteria. Nucleic Acids Res, 2025. 53(2).

16. Michalodimitrakis, K. and M. Isalan, Engineering prokaryotic gene circuits. FEMS Microbiol Rev, 2009. 33(1): p. 27–37.

17. Volk, M.J., et al., Metabolic Engineering: Methodologies and Applications. Chem Rev, 2023. 123(9): p. 5521–5570.

18. Choi, E.N., et al., Expansion of growth substrate range in Pseudomonas putida F1 by mutations in both cymR and todS, which recruit a ring-fission hydrolase CmtE and induce the tod catabolic operon, respectively. Microbiology, 2003. 149(3): p. 795-805.

19. Eaton, R.W., p-Cumate catabolic pathway in Pseudomonas putida Fl: cloning and characterization of DNA carrying the cmt operon. Journal of Bacteriology, 1996. 178(5): p. 1351–1362.

20. Eaton, R.W., p-Cymene catabolic pathway in Pseudomonas putida F1: cloning and characterization of DNA encoding conversion of p-cymene to p-cumate. J Bacteriol, 1997. 179(10): p. 3171–80.

21. Choi, Y.J., et al., Novel, versatile, and tightly regulated expression system for Escherichia coli strains. Appl Environ Microbiol, 2010. 76(15): p. 5058–66.

22. Chang, C., M.D. Phan, and M.A. Schembri, Modified Tn7 transposon vectors for controlled chromosomal gene expression. Appl Environ Microbiol, 2024. 90(10): p. e0155624.

23. Klotz, A., A. Kaczmarczyk, and U. Jenal, A Synthetic Cumate-Inducible Promoter for Graded and Homogenous Gene Expression in Pseudomonas aeruginosa. Appl Environ Microbiol, 2023. 89(6): p. e0021123.

24. Horbal, L., V. Fedorenko, and A. Luzhetskyy, Novel and tightly regulated resorcinol and cumate-inducible expression systems for Streptomyces and other actinobacteria. Appl Microbiol Biotechnol, 2014. 98(20): p. 8641–55.

25. Kaczmarczyk, A., J.A. Vorholt, and A. Francez-Charlot, Cumate-inducible gene expression system for sphingomonads and other Alphaproteobacteria. Appl Environ Microbiol, 2013. 79(21): p. 6795–802.

26. Mullick, A., et al., The cumate gene-switch: a system for regulated expression in mammalian cells. BMC Biotechnol, 2006. 6: p. 43.

27. Schuster, L.A. and C.R. Reisch, A plasmid toolbox for controlled gene expression across the Proteobacteria. Nucleic Acids Res, 2021. 49(12): p. 7189–7202.

28. Venkataraman, M., et al., Synthetic Biology Toolbox for Nitrogen-Fixing Soil Microbes. ACS Synth Biol, 2023. 12(12): p. 3623–3634.

29. Bzdyl, N.M., et al., Pathogenicity and virulence of Burkholderia pseudomallei. Virulence, 2022. 13(1): p. 1945–1965.

30. Mahenthiralingam, E. and P. Vandamme, Taxonomy and pathogenesis of the Burkholderia cepacia complex. Chron Respir Dis, 2005. 2(4): p. 209–17.

31. Loutet, S.A. and Valvano M.A., A decade of Burkholderia cenocepacia virulence determinant research. Infect Immun, 2010. 78(10): p. 4088–100.

32. Lamothe, J., et al., Intracellular survival of Burkholderia cenocepacia in macrophages is associated with a delay in the maturation of bacteria-containing vacuoles. Cell Microbiol, 2007. 9(1): p. 40–53.

33. Flannagan, R.S., T. Linn, and M.A. Valvano, A system for the construction of targeted unmarked gene deletions in the genus Burkholderia. Environ Microbiol, 2008. 10(6): p. 1652–60.

34. Aubert, D.F., M.A. Hamad, and M.A. Valvano, A markerless deletion method for genetic manipulation of Burkholderia cenocepacia and other multidrug-resistant gram-negative bacteria. Methods Mol Biol, 2014. 1197: p. 311–27.

35. Lefebre, M.D. and M.A. Valvano, Construction and evaluation of plasmid vectors optimized for constitutive and regulated gene expression in Burkholderia cepacia complex isolates. Appl Environ Microbiol, 2002. 68(12): p. 5956–64.

36. Cardona, S.T. and M.A. Valvano, An expression vector containing a rhamnose-inducible promoter provides tightly regulated gene expression in Burkholderia cenocepacia. Plasmid, 2005. 54(3): p. 219–28.

37. Qiu, D., et al., PBAD-based shuttle vectors for functional analysis of toxic and highly regulated genes in Pseudomonas and Burkholderia spp. and other bacteria. Appl Environ Microbiol, 2008. 74(23): p. 7422–6.

38. Siegele, D.A. and J.C. Hu, Gene expression from plasmids containing the araBAD promoter at subsaturating inducer concentrations represents mixed populations. Proc Natl Acad Sci U S A, 1997. 94(15): p. 8168–72.

39. Yap, Z.L., et al., A CRISPR-Cas-associated transposon system for genome editing in Burkholderia cepacia complex species. Appl Environ Microbiol, 2024. 90(7): p. e0069924.

40. Hogan, A.M., et al., A Broad-Host-Range CRISPRi Toolkit for Silencing Gene Expression in Burkholderia. ACS Synth Biol, 2019. 8(10): p. 2372–2384.

41. Bloodworth, R.A., A.S. Gislason, and S.T. Cardona, Burkholderia cenocepacia conditional growth mutant library created by random promoter replacement of essential genes. Microbiologyopen, 2013. 2(2): p. 243–58.

42. Hogan, A.M., et al., Improved Dynamic Range of a Rhamnose-Inducible Promoter for Gene Expression in Burkholderia spp. Appl Environ Microbiol, 2021. 87(18): p. e0064721.

43. Lewis, J.M. and N.E. Scott, CRISPRi-Mediated Silencing of Burkholderia O-Linked Glycosylation Systems Enables the Depletion of Glycosylation Yet Results in Modest Proteome Impacts. Journal of Proteome Research, 2023. 22(6): p. 1762–1778.

44. Lithgow, K.V., et al., A general protein O-glycosylation system within the Burkholderia cepacia complex is involved in motility and virulence. Mol Microbiol, 2014. 92(1): p. 116–37.

45. Oppy, C.C., et al., Loss of O-Linked Protein Glycosylation in Burkholderia cenocepacia Impairs Biofilm Formation and Siderophore Activity and Alters Transcriptional Regulators. mSphere, 2019. 4(6).

46. Hoang, T.T., et al., Integration-proficient plasmids for Pseudomonas aeruginosa: site-specific integration and use for engineering of reporter and expression strains. Plasmid, 2000. 43(1): p. 59–72.

47. Hamad Mohamad, A., M. Skeldon Alexander, and A. Valvano Miguel, Construction of Aminoglycoside-Sensitive Burkholderia cenocepacia Strains for Use in Studies of Intracellular Bacteria with the Gentamicin Protection Assay. Applied and Environmental Microbiology, 2010. 76(10): p. 3170–3176.

48. Hanko, E.K.R., et al., A genome-wide approach for identification and characterisation of metabolite-inducible systems. Nature Communications, 2020. 11(1): p. 1213.

49. Hamlin, J.N.R., R.A.M. Bloodworth, and S.T. Cardona, Regulation of phenylacetic acid degradation genes of Burkholderia cenocepacia K56-2. BMC Microbiology, 2009. 9(1): p. 222.

50. Chakravarthy, S., B. Huot, and B.H. Kvitko, Effector Translocation: Cya Reporter Assay. Methods Mol Biol, 2017. 1615: p. 473-487.

51. Mills, E., et al., Real-time analysis of effector translocation by the type III secretion system of enteropathogenic Escherichia coli. Cell Host Microbe, 2008. 3(2): p. 104–13.

52. Sonntag, F., et al., Engineering Methylobacterium extorquens for de novo synthesis of the sesquiterpenoid α-humulene from methanol. Metabolic Engineering, 2015. 32: p. 82–94.

53. Pöschel, L., et al., Expression of toxic genes in Methylorubrum extorquens with a tightly repressed, cumate-inducible promoter. Antonie van Leeuwenhoek, 2023. 116(12): p. 1285–1294.

54. Agulló, L., et al., p-Cymene Promotes Its Catabolism through the p-Cymene and the p-Cumate Pathways, Activates a Stress Response and Reduces the Biofilm Formation in Burkholderia xenovorans LB400. PLoS One, 2017. **12**(1): p. e0169544.

55. Morya, R., D. Salvachúa, and I.S. Thakur, Burkholderia: An Untapped but Promising Bacterial Genus for the Conversion of Aromatic Compounds. Trends in Biotechnology, 2020. 38(9): p. 963–975.

56. Xu, X. and M. Hensel, Systematic analysis of the SsrAB virulon of Salmonella enterica. Infect Immun, 2010. 78(1): p. 49–58.

57. Xu, X., et al., Efficacy of intracellular activated promoters for generation of Salmonella-based vaccines. Infect Immun, 2010. 78(11): p. 4828–38.

58. Lewis, J.M., et al., Glycoproteomic and proteomic analysis of Burkholderia cenocepacia reveals glycosylation events within FliF and MotB are dispensable for motility. Microbiol Spectr, 2024. 12(6): p. e0034624.

59. Hayes, A.J., et al., Burkholderia PglL enzymes are Serine preferring oligosaccharyltransferases which target conserved proteins across the Burkholderia genus. Communications Biology, 2021. 4(1): p. 1045.

60. Paszti, S., et al., Unraveling Burkholderia cenocepacia H111 fitness determinants using two animal models. mSystems, 2025. 10(4): p. e0135424.

61. Fathy Mohamed, Y., et al., A general protein O-glycosylation machinery conserved in Burkholderia species improves bacterial fitness and elicits glycan immunogenicity in humans. J Biol Chem, 2019. 294(36): p. 13248–13268.

62. Garcia, E.C., et al., Burkholderia BcpA mediates biofilm formation independently of interbacterial contact-dependent growth inhibition. Mol Microbiol, 2013. 89(6): p. 1213–25.

63. Gibson, D.G., et al., Enzymatic assembly of DNA molecules up to several hundred kilobases. Nat Methods, 2009. 6(5): p. 343–5.

64. Schindelin, J., et al., Fiji: an open-source platform for biological-image analysis. Nat Methods, 2012. 9(7): p. 676–82.

65. Rappsilber, J., Y. Ishihama, and M. Mann, Stop and go extraction tips for matrix-assisted laser desorption/ionization, nanoelectrospray, and LC/MS sample pretreatment in proteomics. Anal Chem, 2003. 75(3): p. 663–70.

66. Rappsilber, J., M. Mann, and Y. Ishihama, Protocol for micro-purification, enrichment, pre-fractionation and storage of peptides for proteomics using StageTips. Nat Protoc, 2007. 2(8): p. 1896–906.

67. Saba, J., et al., Increasing the productivity of glycopeptides analysis by using higher-energy collision dissociation-accurate mass-product-dependent electron transfer dissociation. Int J Proteomics, 2012. 2012: p. 560391.

68. Caval, T., J. Zhu, and A.J.R. Heck, Simply Extending the Mass Range in Electron Transfer Higher Energy Collisional Dissociation Increases Confidence in N- Glycopeptide Identification. Anal Chem, 2019. 91(16): p. 10401–10406.

69. Polasky, D.A., et al., Multi-attribute Glycan Identification and FDR Control for Glycoproteomics. Mol Cell Proteomics, 2022: p. 100205.

70. Kong, A.T., et al., MSFragger: ultrafast and comprehensive peptide identification in mass spectrometry-based proteomics. Nat Methods, 2017. 14(5): p. 513–520.

71. Polasky, D.A., et al., Fast and comprehensive N- and O-glycoproteomics analysis with MSFragger-Glyco. Nat Methods, 2020. 17(11): p. 1125–1132.

72. Holden, M.T., et al., The genome of Burkholderia cenocepacia J2315, an epidemic pathogen of cystic fibrosis patients. J Bacteriol, 2009. 191(1): p. 261–77.

73. Cox, J., et al., Accurate proteome-wide label-free quantification by delayed normalization and maximal peptide ratio extraction, termed MaxLFQ. Mol Cell Proteomics, 2014. 13(9): p. 2513–26.

74. Tyanova, S., et al., The Perseus computational platform for comprehensive analysis of (prote)omics data. Nat Methods, 2016. 13(9): p. 731–40.

75. Wickham, H., *ggplot2: Elegant Graphics for Data Analysis*. 2016: Springer-Verlag New York.

76. Demichev, V., et al., DIA-NN: neural networks and interference correction enable deep proteome coverage in high throughput. Nat Methods, 2020. 17(1): p. 41–44.

77. Keilhauer, E.C., M.Y. Hein, and M. Mann, Accurate protein complex retrieval by affinity enrichment mass spectrometry (AE-MS) rather than affinity purification mass spectrometry (AP-MS). Mol Cell Proteomics, 2015. 14(1): p. 120–35.

